# Benchmarking of bioinformatics tools for the hybrid *de novo* assembly of human whole-genome sequencing data

**DOI:** 10.1101/2024.05.28.595812

**Authors:** Adrián Muñoz-Barrera, Luis A. Rubio-Rodríguez, David Jáspez, Almudena Corrales, Itahisa Marcelino-Rodriguez, José M. Lorenzo-Salazar, Rafaela González-Montelongo, Carlos Flores

## Abstract

Accurate and complete *de novo* assembled genomes sustain variant identification and catalyze the discovery of new genomic features and biological functions. However, accurate and precise *de novo* assemblies of large and complex genomes remains a challenging task. Long-read sequencing data alone or in hybrid mode combined with more accurate short-read sequences facilitate the *de novo* assembly of genomes. A number of software exists for *de novo* genome assembly from long-read data although specific performance comparisons to assembly human genomes are lacking. Here we benchmarked 11 different pipelines including four long-read only assemblers and three hybrid assemblers, combined with four polishing schemes for *de novo* genome assembly of a human reference material sequenced with Oxford Nanopore Technologies and Illumina. In addition, the best performing choice was validated in a non-reference routine laboratory sample. Software performance was evaluated by assessing the quality of the assemblies with QUAST, BUSCO, and Merqury metrics, and the computational costs associated with each of the pipelines were also assessed. We found that Flye was superior to all other assemblers, especially when relying on Ratatosk error-corrected long-reads. Polishing improved the accuracy and continuity of the assemblies and the combination of two rounds of Racon and Pilon achieved the best results. The assembly of the non-reference sample showed comparable assembly metrics as those of the reference material. Based on the results, a complete optimal analysis pipeline for the assembly, polishing, and contig curation developed on Nextflow is provided to enable efficient parallelization and built-in dependency management to further advance in the generation of high-quality and chromosome-level human assemblies.

## Introduction

Next-generation sequencing technologies (NGS) have enabled to rapidly expand our knowledge of the human genome with unprecedented precision (Goodwin, McPherson, and McCombie 2016). As the Precision Medicine paradigm shift is embraced, the accurate reconstruction of individual human genomes is key, necessitating the deployment of robust bioinformatics tools for *de novo* assembly (Wee et al. 2019) to provide a comprehensive and unbiased understanding of a patient’s DNA sequence. *De novo* genome assembly is crucial for unveiling the full spectrum of genetic diversity among individuals, shedding light on population-specific variation (Kim et al. 2019; Nagasaki et al. 2019; Chao et al. 2023) and rare alleles that may influence disease risk and treatment response (Helal et al. 2022; Deng et al. 2022), among other applications. Third-generation sequencing (TGS), as seen in Oxford Nanopore Technologies (ONT) and Pacific Biosciences (PacBio), relies on long-read capabilities that boost the possibilities of a comprehensive view of the genome (Athanasopoulou et al. 2021). Traditional short-read NGS retains optimal turnaround time and cost-effectiveness. However, the existing algorithms struggle to provide a highly continuous *de novo* genome assembly based on short reads, failing to resolve complex genomic regions, repetitive elements, and most of the structural variants. TGS greatly expands the possibilities (Marx 2021) despite its lower per base accuracy compared to short-read NGS (Perešíni et al. 2021) since a significant portion of the human genome consists of repetitive sequences essential for understanding the genetic basis of diseases (Wagner et al. 2022).

Unlike others, ONT reads typically span thousands of bases (Magi et al. 2018) providing a more comprehensive and contiguous view of genomic regions that are otherwise challenging to resolve (Logsdon, Vollger, and Eichler 2020; Amarasinghe et al. 2020). To harness the benefits of ONT long-reads while addressing their inherent error profiles, two primary strategies are typically employed. One involves error correction of long reads before the actual genome assembly, which is achieved with high coverage sequencing (Salmela et al. 2017; Dohm et al. 2020; Tang et al. 2023). For this, information from multiple reads covering the same genomic region is leveraged to identify and rectify random errors, thereby enhancing the reliability of the raw long-read data. Another strategy polishes the draft assembled sequence obtained from long reads (Lee et al. 2021), so that the draft genome sequence undergoes iterative refinement through alignment with high-quality short-read data or consensus sequences. This step helps to rectify any remaining errors, fine-tuning the accuracy and completeness of the assembled genome. The combination of pre-assembly error correction and post-assembly polishing ensures a more robust and accurate representation of the genome (Y. Chen et al. 2021; Morisse et al. 2021; Fang and Wang 2022).

Despite the importance and the existence of different bioinformatics tools for *de novo* assembly of genomes, there is a lack of studies assessing the benefits and limitations of the available tools for ONT read data, especially in the context of humans (T. Zhang et al. 2022). Here we aimed to benchmark alternative *de novo* genome assembly bioinformatics tools of nanopore data from human whole-genomes. For that, we used a human reference material sequenced with ONT and Illumina, and then validated the performance of the best benchmarked tool in a non-reference routine laboratory sample with lower integrity. We also contribute with a complete best-performing analysis pipeline for assembly, polishing, and contig curation developed on Nextflow enabling parallelization and built-in dependency management.

## Materials and methods

In brief, the study workflow involved the use of data from a human reference material sequenced with ONT and Illumina to benchmark 11 *de novo* and hybrid assembly pipelines, combined with different polishing schemes to improve the accuracy and continuity of the assemblies. The best assembler and polishing scheme were then validated to assemble the complete genome of a non-reference routine human sample with lower integrity.

### Human whole-genome sequence datasets

#### Human data for benchmarking of the assembly pipelines and polishing schemes

For the evaluation of all selected assembly and polishing tools, data from the HG002 sample (or NA24385, Son of Ashkenazi Jewish ancestry) was selected as reference since it is commonly used for calibration, development of genome assembly methods, and laboratory performance measurements as part of the Genome in a Bottle (GIAB) Consortium. The raw Illumina and ONT sequence data from this sample is publicly available (Zook et al. 2016). Briefly, the Illumina dataset was obtained by sequencing on a NovaSeq 6000 System (Illumina, Inc.) and then downsampled to 35X, while the ONT dataset was obtained with PromethION (Oxford Nanopore Technologies) with R9.4 flow cells and base calling was performed using Guppy (v3.6.0), obtaining a genome coverage of 47X (Olson et al. 2022). Note that the mitogenome scaffold was previously reconstructed using an in-house pipeline described elsewhere (García-Olivares et al. 2021) and not assessed here.

#### Human data for validation of the best performing pipeline

To validate the performance of the best pipeline resulting from the benchmarking, we used human whole-genome data from a routine sample from our laboratory (CAN0003). This sample was included in the reference genetic catalog of the Canary Islands population (CIRdb) that has been described elsewhere (Díaz-de Usera et al. 2022). The details for DNA isolation, library preparation, and sequencing were previously described (García-Olivares et al. 2021). Briefly, the short-read dataset was obtained using the Nextera DNA Library Preparation Kit and the sequence was obtained on a HiSeq 4000 Sequencing System (Illumina, Inc.) at the Instituto Tecnológico y de Energías Renovables (ITER, Santa Cruz de Tenerife, Spain). Raw BCL files were demultiplexed and converted to FASTQ files by means of bcl2fastq (v2.20). The ONT long-read dataset was obtained at Keygene (Wageningen, The Netherlands) using the ligation library preparation kit (SQK_LSK109) and sequenced on a PromethION platform (Oxford Nanopore Technologies) with a R9.4.1 flow cell (FLO_PR002) and MinKNOW (v1.14.2) software. After the run, base calling was performed with the Guppy (v5.0.7) neural network based tool. The mitogenome was obtained with the same methods as for the HG002 sample and are not assessed here.

### Bioinformatics workflows

#### Overview

The evaluation of long-read and hybrid *de novo* assembly bioinformatics tools and the polishing pipelines were implemented using command-line interface Bash scripts (**Figure 1A**). Based on the results, the complete analysis pipeline for the assembly, polishing, and contig curation using the best resulting tools (**Figure 1B**) was developed on Nextflow (v23.04.1) (Di Tommaso et al. 2017) workflow management language using the templates and following the best practice guidelines provided by nf-core community (Ewels et al. 2020). This implementation enables efficient parallelization and built-in dependency management through Docker (Merkel 2014) containers and Conda (“Anaconda Software Distribution” 2020) environments. Detailed usage of each bioinformatics tool and the complete pipeline is described in a dedicated repository: https://github.com/genomicsITER/hybridassembly.

**Figure 1.**
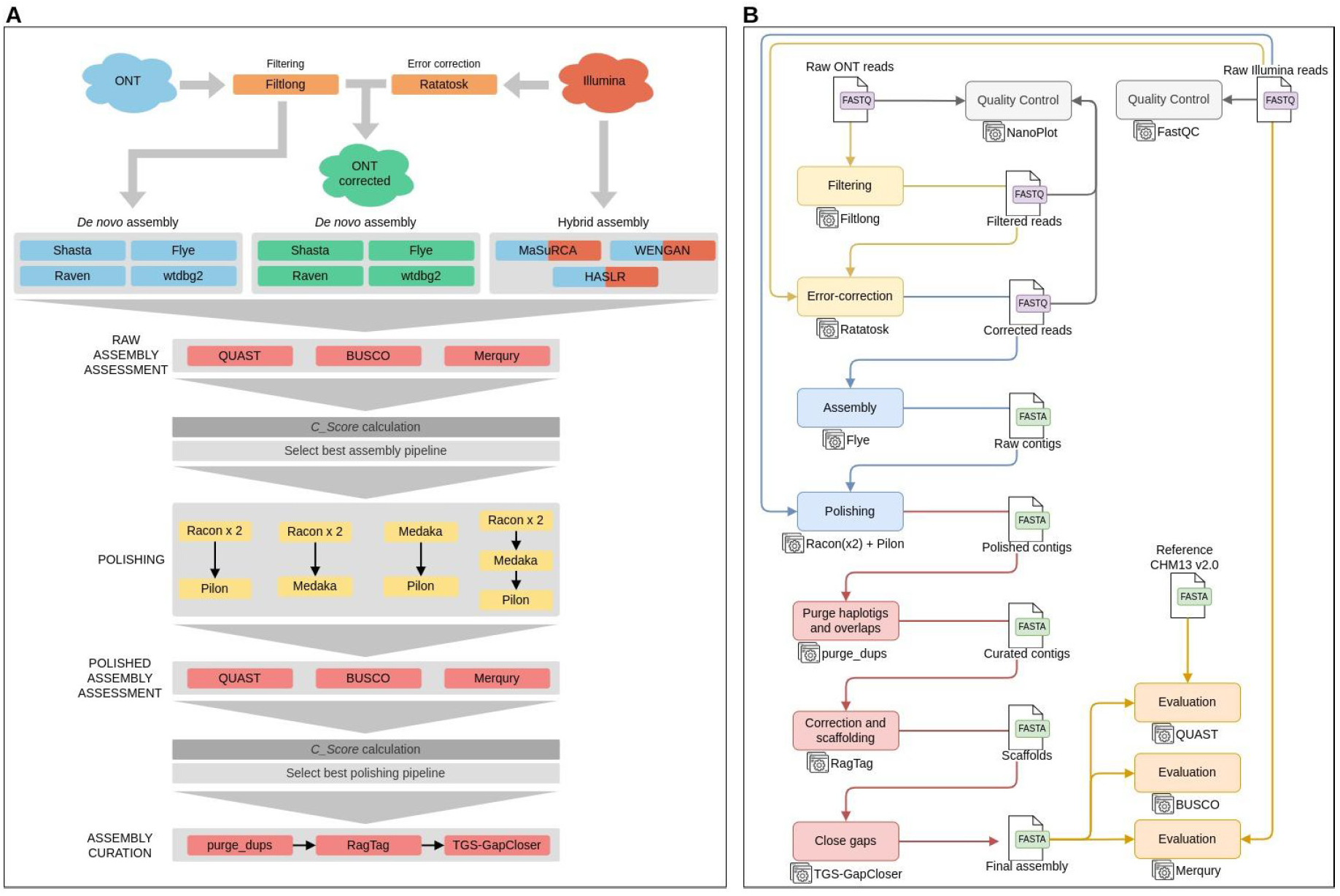
A. Long-read and hybrid *de novo* genome assembly pipelines for evaluation and benchmarking purposes using the reference sample data (HG002) as input. The datasets used in each pipeline were represented by colors: Blue, long-reads from ONT; Red, short-reads from Illumina; Green, long-reads from ONT corrected with short-reads. **B.** Detailed pipeline implemented in Nextflow using best assembly

#### Filtering, error correction, and initial quality control steps

First, Filtlong (v0.2.1) (https://github.com/rrwick/Filtlong) was used to remove long-read sequences shorter than 1,000 bp length from the ONT dataset. A base level error-correction step was performed afterwards using Ratatosk (v0.9.0) (Holley et al. 2021) to correct previous filtered long-reads using Illumina short-reads from the same sample.

Quality control assessments of ONT reads for the reference and validation sample datasets were performed using NanoPlot (v1.39.0) (De Coster et al. 2018) before and after filtering and error correction processes. FastQC (v0.12.1) (Andrews and Others 2017) was used to evaluate raw short-read sequence data.

#### *De novo* genome assemblers benchmarked in this study

We evaluated the performance of four bioinformatics tools for *de novo* genome assembly of long and complex genomes using long-reads from ONT: Shasta (v0.9.0) (Shafin et al. 2020), Flye (v2.9) (Freire, Ladra, and Parama 2021), Raven (v1.8.1) (Vaser and Šikić 2021), and wtdbg2 (v2.5) (Ruan and Li 2019). For these, two datasets of the reference sample (HG002) were used as input: i) the filtered long-reads, and ii) the filtered and the Ratatosk-corrected long-reads. With this, it was possible to evaluate the performance of the *de novo* assembly tools using error-correction processes prior to the assembly steps. For simplicity, we will refer to the first set of assemblies as Shasta, Flye, Raven, and wtdbg2, and to the second set as Corrected_Shasta, Corrected_Flye, Corrected_Raven, and Corrected_wtdbg2. All these assembly tools were run using default parameters.

We also evaluated the performance of the following three hybrid *de novo* assemblers: MaSuRCA (v4.0.8) (Zimin et al. 2013), WENGAN (v0.2) (Di Genova et al. 2020), and HASLR (v0.8a1) (Haghshenas et al. 2020). For these, we used the reference sample dataset combining Illumina short-reads with the filtered ONT reads, and used default software parameters.

#### Assessment of raw genome assemblies

The quality of all resulting assemblies generated by the combination of assemblers and polishing tools was evaluated by means of QUAST (v5.0.2) (Gurevich et al. 2013; Mikheenko et al. 2018) using T2T-CHM13v2.0 as reference genome. BUSCO (v5.3.2) (Simão et al. 2015; Seppey, Manni, and Zdobnov 2019) was used to evaluate gene completeness of assemblies, while Merqury (v1.3) (Rhie et al. 2020) was used as a reference-free assessment tool, to determine their quality using short-read sequencing data.

Inspired by a previous study (X. Zhang et al. 2022), we modified the Comprehensive Score (CS) as an integrative score aggregating several metrics provided by QUAST, BUSCO, and Merqury (further details in the **Supplementary Material**). Briefly, CS integrated six metrics: the number of contigs, N50 in Mbp, number of mismatches and indels per 100 kbp, number of complete genes (completeness) annotated by BUSCO, and the consensus quality value (QV) estimated by Merqury.

#### Comparing alternative polishing tools in the raw assembly with the best CS

The combination of input datasets, preprocessing steps, and assembly pipelines provided several possibilities. To simplify the comparisons of alternative polishing tools, we applied four schemes combining alternative state-of-the-art polishing tools, Racon (v1.5.0) (Vaser et al. 2017), Medaka (v1.6.0) (https://community.nanoporetech.com), and Pilon (v1.24) (Walker et al. 2014), on the assembly with the best CS obtained in the previous stage.

Four different polishing schemes were tested in order to assess the impact of running multiple rounds of polishing using only long reads or using both long and short reads (see **Figure 1A**): a) two rounds of Racon combined with Pilon (referred as Racon_Pilon), b) two rounds of Racon combined with Medaka (referred as Racon_Medaka), c) one round of each Medaka combined with Pilon (referred as Medaka_Pilon), and d) all three combined with two rounds of Racon followed by one round each of Medaka and Pilon (referred as Racon_Medaka_Pilon).

Results of these polishing schemes were evaluated as for the raw assemblies, i.e., using QUAST, BUSCO, and Merqury to calculate the CS of the polished assemblies.

#### Contig curation, scaffolding, and gap-filling

Polished contigs resulting from the polishing schema with best CS were curated using purge_dups (v1.2.6) (Guan et al. 2019) to remove haplotigs and contig overlaps based on read depth, reducing heterozygous duplication and increasing assembly continuity while maintaining completeness of the primary assembly. Potential misassemblies of curated contigs were further corrected and then ordered and oriented in the scaffolding step by means of RagTag (v2.1.0) (Alonge et al. 2022) using the T2T-CHM13v2.0 as the reference genome. Gaps (Ns) present in the scaffolds were attempted to be filled using TGS-GapCloser (v1.2.1) (Xu et al. 2020).

#### Validation of the pipeline with best results in a dataset from a routine sample

Finally, in order to assess the performance of these tools in a real case scenario, we used data from a routine laboratory sample with lower integrity than the reference material. The data from the validation sample was processed using the best assembly pipeline and polishing scheme, including curation, scaffolding, and closing-gaps steps (**Figure 1B**). The final assembly was assessed and compared against the final assembly of HG002 with QUAST, BUSCO, and Merqury.

### Computational time and memory usage

Some of the key aspects to take into account in the context of this benchmarking are the computational time and resources associated with the *de novo* genome assembly process (Kleftogiannis, Kalnis, and Bajic 2013). The performance of each selected assembly tool and polishing pipelines in this study were evaluated in terms of computational efficiency and time consumption by monitoring each process to obtain the total time execution and the peak of memory consumption.

### Hardware resources

All the bioinformatics processes were conducted in two settings: an HPC cluster infrastructure, namely the TeideHPC (described here: https://teidehpc.iter.es), and a local workstation running CentOS 7 with 2 Intel® Xeon® Platinum 8358 CPUs at 2.60 GHz and with 2 TB of RAM.

## Results

### Initial quality control of the reference HG002 sample dataset

The Illumina HG002 dataset consists of 415 Mreads with 151 bp length, providing a genome coverage of 39X. The raw ONT HG002 dataset consists of 19.3 Mreads with a N50 value of 50.3 kbp and a mean read quality of 8 in the Phred scale (**Table 1**).

**Table 1.**
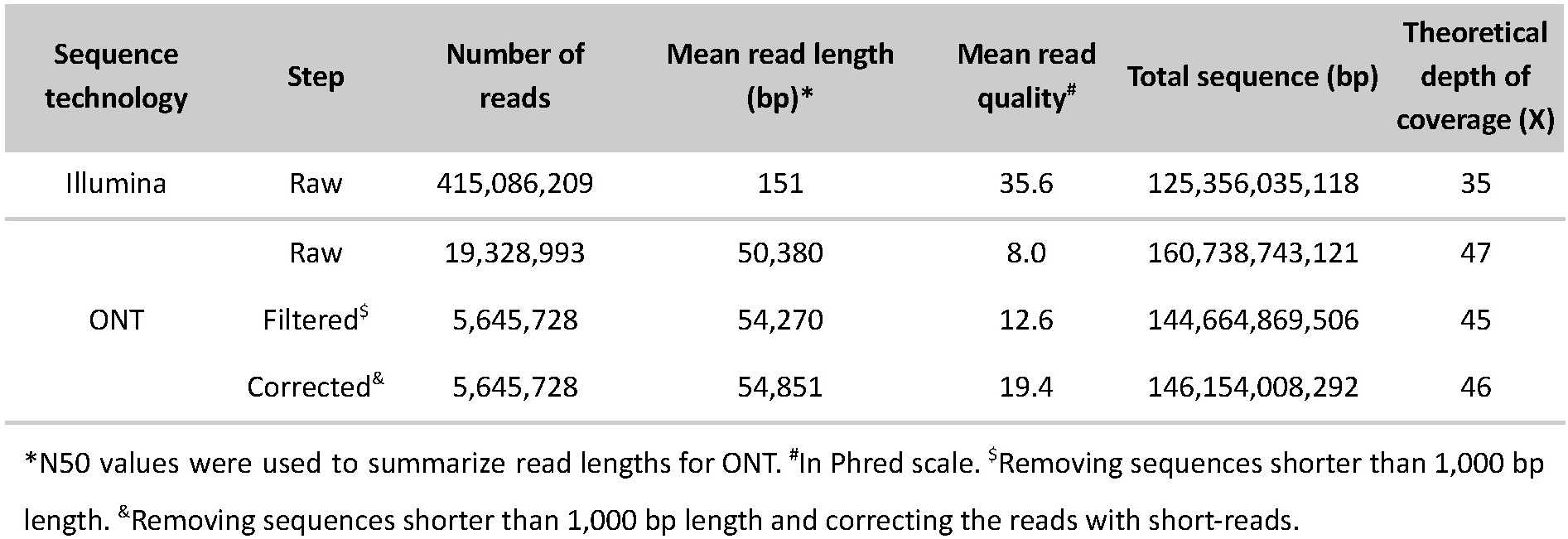
Raw sequence and preprocessed data characteristics of the HG002 genome.

Although filtering out shorter reads implies a drastic reduction in the total number of reads (**Table 1**), the resulting dataset maintains 90% of the represented sequences, raising the N50 from 50.3 kbp to 54.3 kbp and increasing the mean read quality from 8.0 to 12.6. With the read error-correction, the N50 value improved slightly, as well as the total the represented sequences and the genome coverage. However, a substantial improvement was observed in terms of read quality.

Both filtered and corrected ONT datasets were used as input data for those assemblers that only use long-reads to investigate the impact of error-correction in the resulting assemblies. For the hybrid assemblers, the filtered ONT and raw Illumina datasets were used as input datasets.

### HG002 assembly results

Using previously preprocessed ONT and Illumina datasets of the reference HG002 sample, a total of 11 pipelines (resulting from the combinations of preprocessing datasets and assembly tools) were benchmarked based on the metrics obtained with QUAST, BUSCO, and Merqury (see **Table 2** and Supplementary Table 2).

**Table 2.**
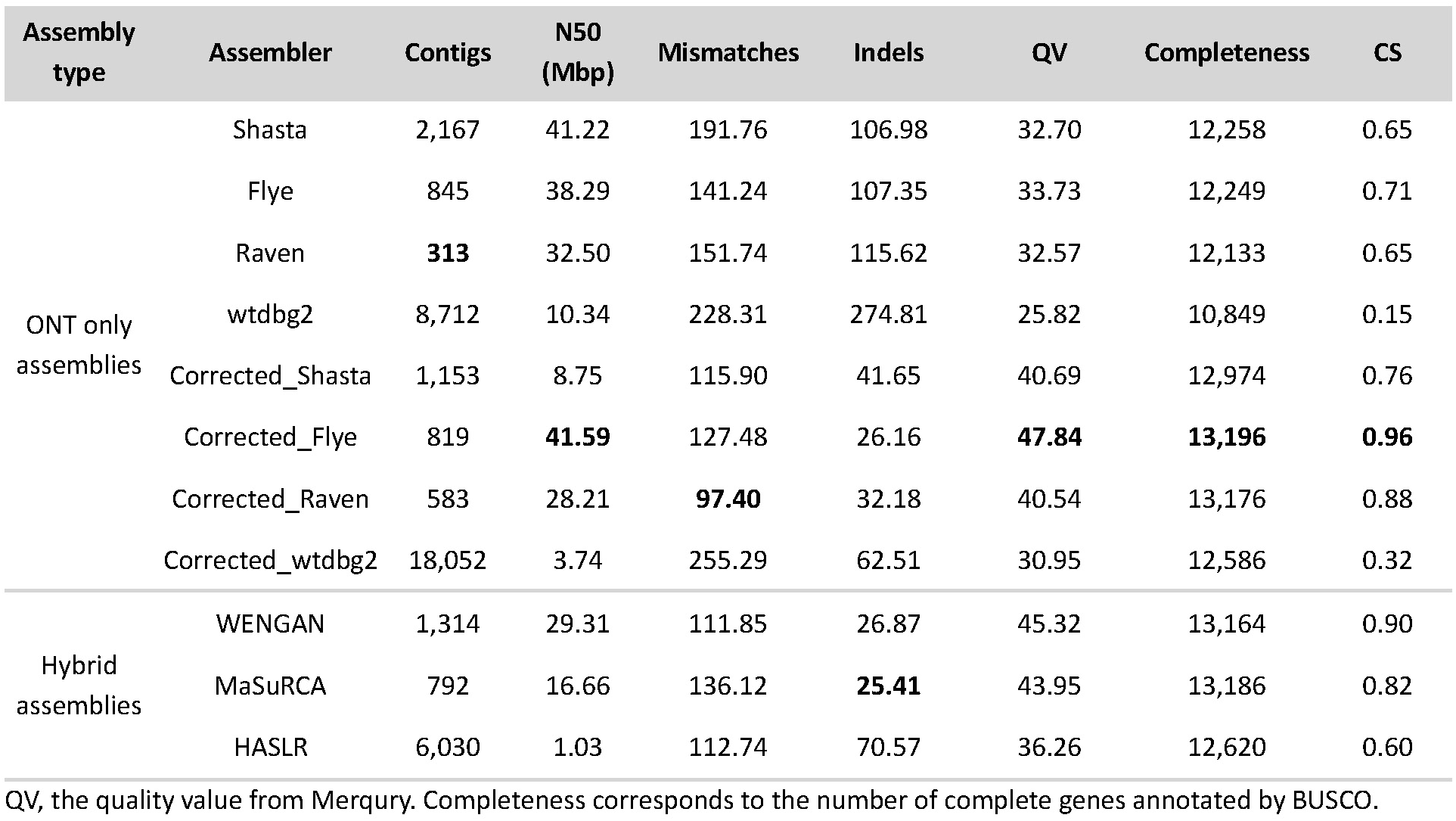
Summary of HG002 assembly results and Comprehensive Scores (CS) obtained for each preprocessing strategy and *de novo* genome assembly pipeline. Contigs, N50 length (Mbp), mismatches (per 100 kbp), and indels (per 100 kbp) were extracted from QUAST using T2T-CHM13v2.0 as reference. The best value for each metric is shown in bold.

| Assembly type | Assembler | Contigs | N50 (Mbp) | Mismatches | Indels | QV | Completeness | CS |
| --- | --- | --- | --- | --- | --- | --- | --- | --- |
| ONT only assemblies | Shasta | 2,167 | 41.22 | 191.76 | 106.98 | 32.70 | 12,258 | 0.65 |
|  | Flye | 845 | 38.29 | 141.24 | 107.35 | 33.73 | 12,249 | 0.71 |
|  | Raven | **313** | 32.50 | 151.74 | 115.62 | 32.57 | 12,133 | 0.65 |
|  | wtdbg2 | 8,712 | 10.34 | 228.31 | 274.81 | 25.82 | 10,849 | 0.15 |
|  | Corrected_Shasta | 1,153 | 8.75 | 115.90 | 41.65 | 40.69 | 12,974 | 0.76 |
|  | Corrected_Flye | 819 | **41.59** | 127.48 | 26.16 | **47.84** | **13,196** | **0.96** |
|  | Corrected_Raven | 583 | 28.21 | **97.40** | 32.18 | 40.54 | 13,176 | 0.88 |
|  | Corrected_wtdbg2 | 18,052 | 3.74 | 255.29 | 62.51 | 30.95 | 12,586 | 0.32 |
| Hybrid assemblies | WENGAN | 1,314 | 29.31 | 111.85 | 26.87 | 45.32 | 13,164 | 0.90 |
|  | MaSuRCA | 792 | 16.66 | 136.12 | **25.41** | 43.95 | 13,186 | 0.82 |
|  | HASLR | 6,030 | 1.03 | 112.74 | 70.57 | 36.26 | 12,620 | 0.60 |

QV, the quality value from Merquy. Completeness corresponds to the number of complete genes annotated by BUSCO.

In terms of contiguity, Flye, Raven, Corrected_Flye, Corrected_Raven, and MaSuRCA had the least number of contigs (<1,000 contigs), wtdbg2 and Corrected_wtdbg2 being the options with more fragmented assemblies (8,712 and 18,052 contigs, respectively). Shasta, Flye, and Corrected_Flye showed the highest N50 values, near 40 Mbp. In contrast, Corrected_Shasta, Corrected_wtdbg2, and HASLR obtained assemblies with N50 values under 10 Mbp. The largest contig was returned by Shasta (138.11 Mbp), followed by Corrected_Flye (109.82 Mbp), WENGAN (109.74 Mbp), Raven (109.44 Mbp), and Flye (108.41 Mbp). Despite this, the retrieved total length of each was similar, ranging from 2.73 Gbp to 2.93 Gbp. The exception was Corrected_wtdbg2, which returned a total length of 3.28 Gbp, possibly due to the extremely high number of contigs returned.

Regarding completeness, Shasta, Flye, and Corrected_Flye showed a NA50 value over 30 Mbp providing a genome fraction above 90%. According to BUSCO, Corrected_Flye had the most complete gene number, closely followed by MaSuRCA, Corrected_Raven, and WENGAN, with more than 95% of completeness reported by BUSCO. Based on Merqury metrics, Corrected_Flye, WENGAN, and MaSuRCA assemblers had the best k-mer completeness (>97%) and Quality Value (QV >43) (**Figure 2**). These results support that the combination with short-read sequencing data, either in the read error correction stage or at the assembly step, improves the *de novo* genome assembly.

**Figure 2.**
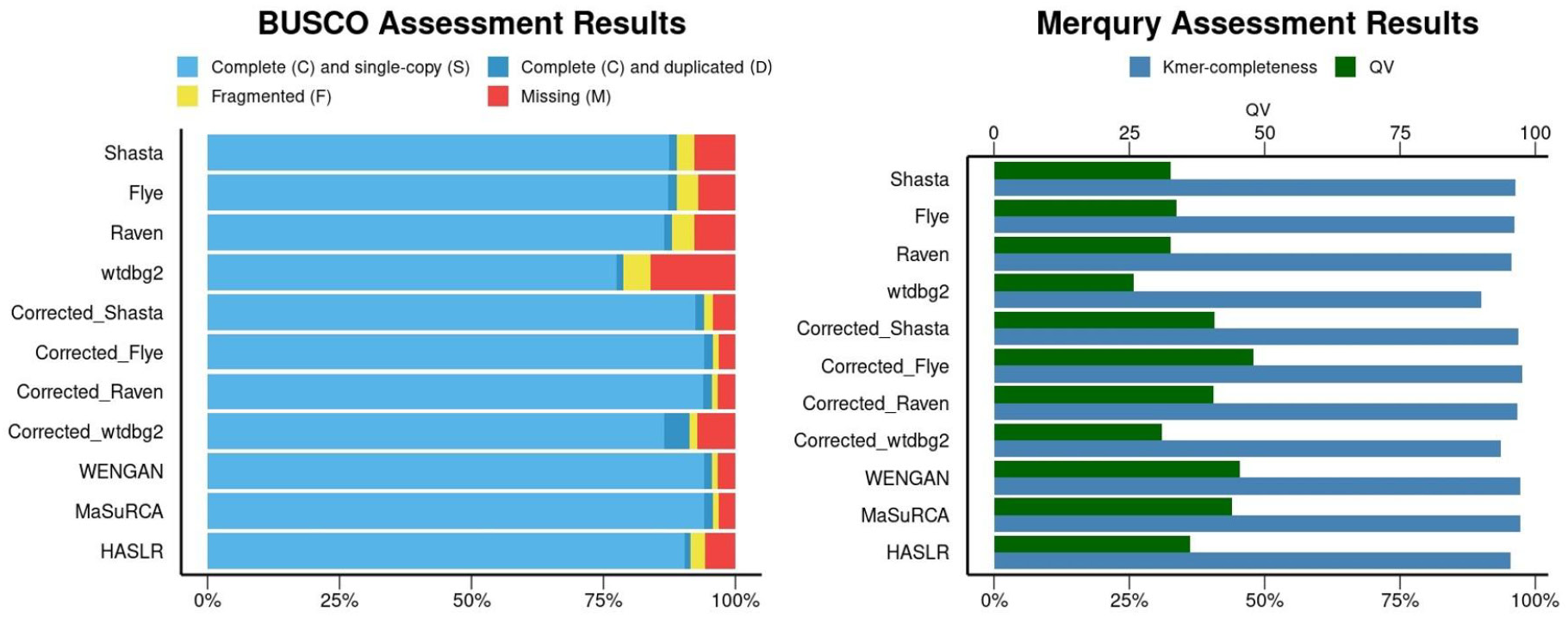
BUSCO and Merqury for HG002 assembly results for all 11 *de novo* genome assembly pipelines.

Based on the correctness or accuracy of the assemblers, the best results were returned by Corrected_Raven, with 97.4 mismatches per 100 kbp, and Corrected_Flye, WENGAN, and MaSuRCA with nearly 25 indels per 100 kbp. The lower number of misassemblies were exhibited by HASLR (113), which also showed a small number of mismatches and indels, although with suboptimal results in terms of contiguity and completeness.

CS calculations for the 11 pipelines showed that Corrected_Flye and WENGAN were superior for *de novo* genome assembly combining ONT and Illumina data. However, Corrected_Flye provided the highest contiguity, resulting from a lower number of contigs (819) and a higher N50 value (41.59 Mbp). Corrected_Flye also had the best accuracy and completeness. Further details of the CS calculations can be found in the **Supplementary Table 3**.

Based on the previous results, the resulting assembly of Corrected_Flye was used in the following steps as the input to evaluate alternative polishing schemes. Overall, the polished assemblies show slight improvements in assembly quality. The schemes that used multiple rounds of Racon as a first polishing step showed greater improvements in contiguity, decreasing the number of contigs from 819 to 800, but maintaining the assembled size and N50. Both, Racon and Medaka introduced more mismatches and indels that were subsequently corrected by the use of Pilon, resulting in a final improvement of the assembly correctness. However, in terms of completeness, the use of these polishing pipelines did not show differences, as the QV metric and the BUSCO completeness remained nearly identical in almost all situations, with the Racon_Medaka scheme being the one showing the worst results (see **Table 3** and **Supplementary Table 4**).

**Table 3.** Summary of HG002 polishing results and CS values obtained for each scheme based on the best assembly (Corrected_Flye). Contigs, N50 length (Mbp), mismatches (per 100kbp), and indels (per 100kbp) were extracted from QUAST using T2T-CHM13v2.0 as reference. The best value of each metric is shown in bold.

| Polishing scheme | Contigs | N50 (Mbp) | Mismatches | Indels | QV | Completeness | CS |
| --- | --- | --- | --- | --- | --- | --- | --- |
| Racon_Pilon | **800** | 41.59 | 131.39 | 24.53 | **46.58** | 13,198 | **0.65** |
| Racon_Medaka | **800** | **41.62** | 133.99 | 28.02 | 42.68 | 13,196 | 0.33 |
| Medaka_Pilon | 819 | 41.59 | **125.82** | **24.21** | 45.77 | **13,199** | 0.63 |
| Racon_Medaka_Pilon | **800** | 41.60 | 127.93 | 24.57 | 44.21 | 13,196 | 0.56 |

QV, the quality value from Merqury. Completeness corresponds to the number of complete genes annotated by BUSCO.

Based on the CS values, the results revealed that Racon_Pilon was the best polishing scheme, closely followed by Medaka_Pilon. Further details underlying CS calculations can be found in **Supplementary Table 5**.

Finally, contig curation, scaffolding, and gap-filling steps were conducted using the polished draft assembly generated by the Racon_Pilon scheme. Resulting metrics from QUAST, BUSCO, and Merqury of the final curated *de novo* genome assembly are shown in **Table 4**. The complete assembly statistics of each curation step are shown in **Supplementary Table 6**.

**Table 4.**
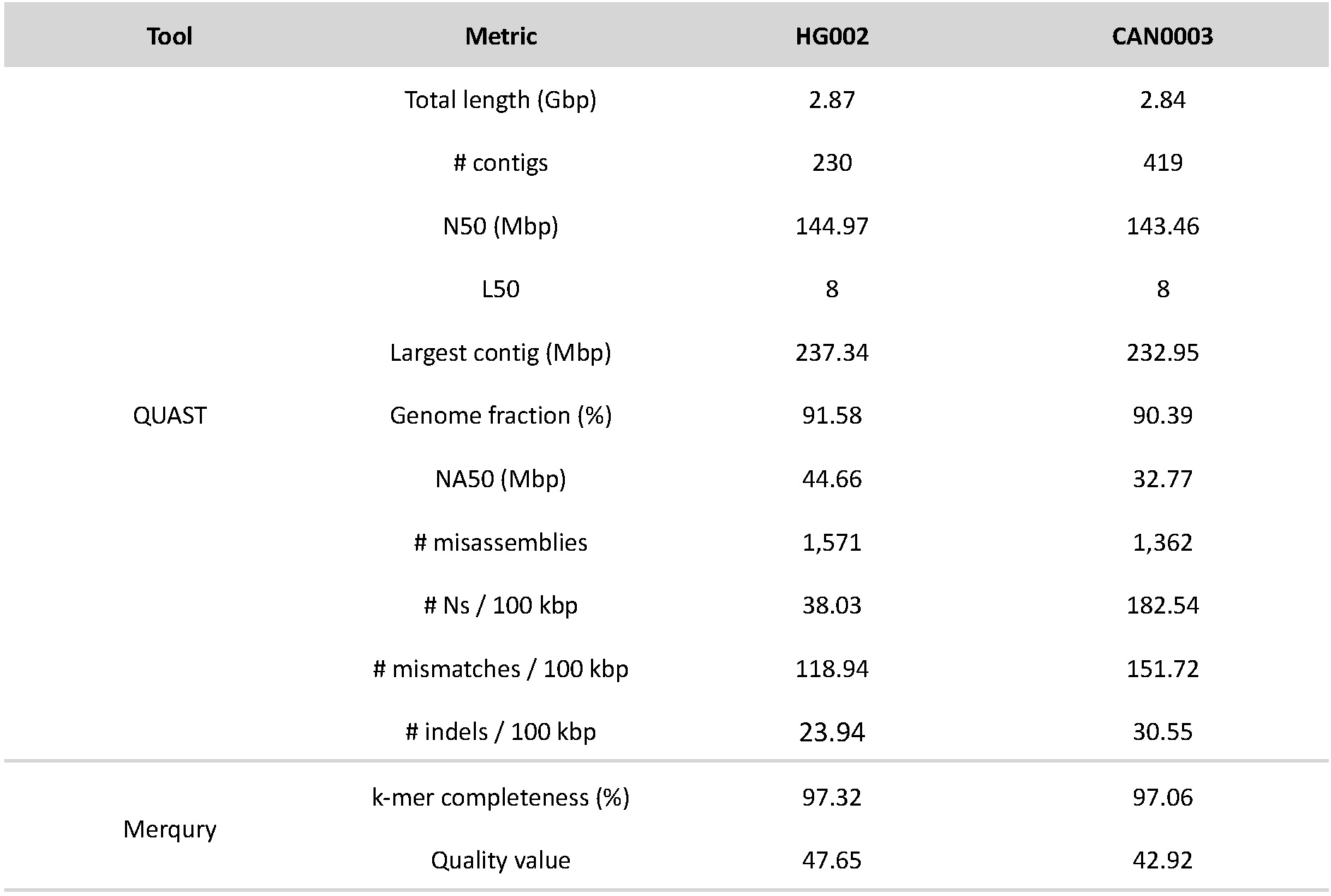
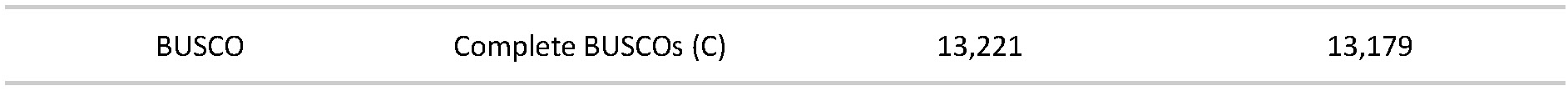
Summary statistics of final *de novo* genome assemblies of HG002 and CAN0003 after contig curation, scaffolding, and gap-filling steps. The T2T-CHM13v2.0 genome was used as the reference in QUAST evaluations.

After contig curation, scaffolding, and gap-filling, the resulting assembly had 25 scaffolds, representing the 22 autosomes, X and Y sexual chromosomes, the mitogenome, and 207 unplaced contigs. Not considering the unplaced contigs, the total length of the assembly was 2,845,020,552 bp, including a total of 183 gaps summing up a total gap length of 1,092,630 bp. A detailed comparison of chromosome lengths and gaps between the assembled HG002 genome and the T2T-CHM13v2.0 reference can be found in the **Supplementary Table 7**.

### Computational time and memory usage

The computational resources required by each assembler in the local workstation setting were diverse, with computational runtimes ranging from 1.53 to 38.6 hours, and a memory usage peak ranging from 107 to 1,471 GB of RAM (**Figure 3**). In terms of computational runtime, Shasta and Corrected_Shasta proved to be the fastest (2.4 and 1.5 h, respectively), although at the cost of a high (>700 GB of RAM) memory usage peak. On the opposite, WENGAN was one of the tools with the highest computational costs both in terms of runtime (23.2 h) and memory usage peak (1,471 GB of RAM). Raven and Corrected_Raven offered a situation of compromise, having a low resource intensity necessitating 107 GB of RAM, albeit still maintaining a reasonably low execution runtimes (10.1 and 9.4 h, respectively).

**Figure 3.**
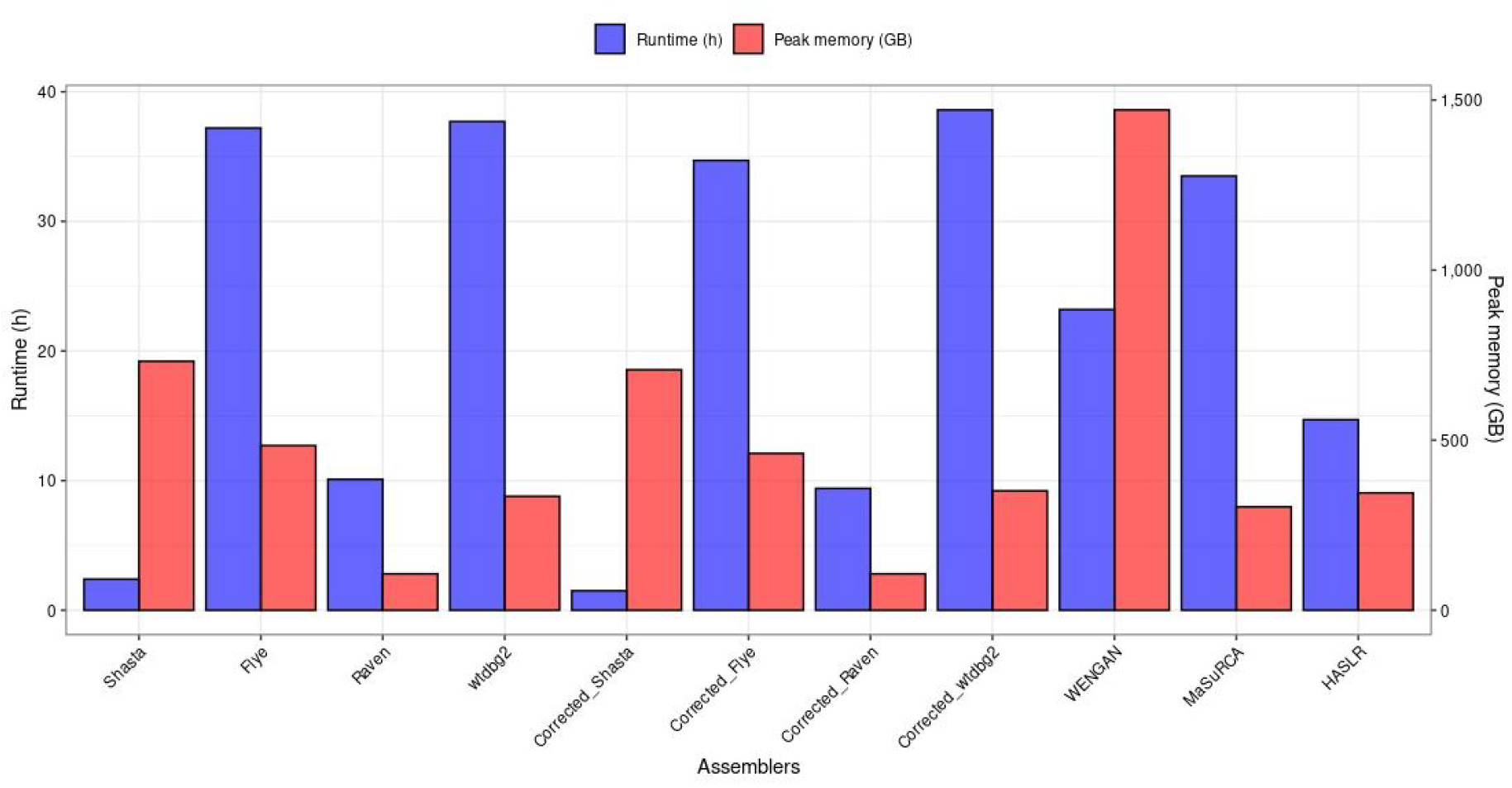
Computational resources (runtime and memory usage peak) required for the *de novo* genome assembly of HG002.

Regarding the polishing schemes, Racon_Pilon and Medaka_Pilon showed similar computational runtimes (42.1 and 41.3 h, respectively). In terms of memory usage, the Racon_Medaka_Pilon strategy, despite its longer computational time (60.4 h) maintains the same memory usage as Racon_Pilon (447 GB) due to the high memory cost of Racon. On the other hand, the Medaka_Pilon scheme has lower memory requirements, showing a peak memory RAM of 365 GB.

### Initial quality control of the validation CAN0003 sample dataset

Multiple sequencing statistics including total sequenced reads and bases, average read length and quality, and the read length N50, were calculated for each dataset of this sample (**Table 5**). The Illumina dataset consisted of 341 Mreads with 151 bp length, providing a theoretical genome coverage of 29X. Raw ONT dataset consisted of 10.9 Mreads with a N50 value of 17.2 kbp and a mean read quality of 10.7. After filtration and correction steps, this dataset resulted in 7.5 Mreads with a N50 value of 18.3 kbp and a mean read quality of 18.4. As expected due to the laboratory handling and storage conditions of routine samples, a notable decrease in the number of reads and their overall shorter lengths were evident in comparison to those of HG002.

**Table 5.**
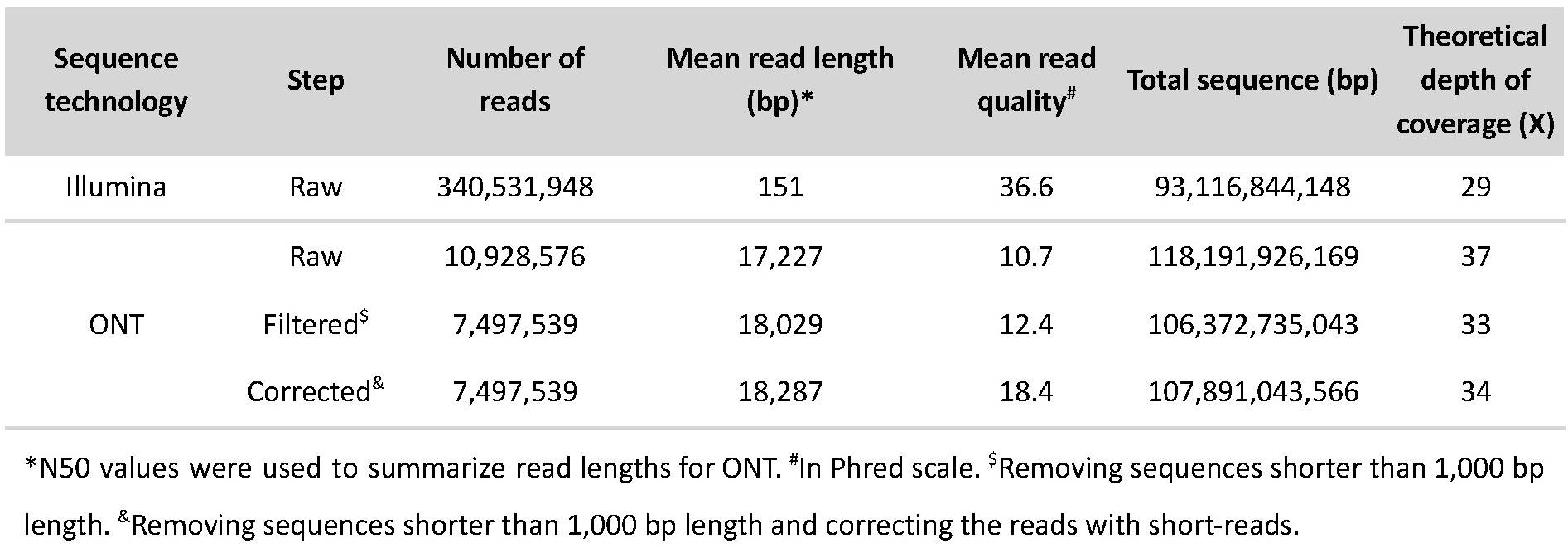
Raw sequence and preprocessed data characteristics of the CAN0003 validation sample.

#### CAN0003 assembly results

Based on the optimal performance on HG002, we used Corrected_Flye for *de novo* genome assembly and Racon_Pilon as the polishing scheme. The assembly results from QUAST, BUSCO, and Merqury of the final curated assembly are shown in **Table 4**. Further details of each curation step are shown in **Supplementary Table 8**. Compared to HG002, results for CAN0003 were similar in terms of contiguity (**Figure 4**), showing more contigs (n=419), although similar N50 values (143.46 Mbp) and a similar total length (2.84 Gbp). In terms of completeness, the results show 3.8 times more Ns/100 kbp (146.60 in contrast to 38.28 of HG002) and a similar number of complete BUSCOs (13,179). As for correctness, the CAN0003 genome assembly was worse than that of HG002, including 151.72 mismatches/100 kbp, and 30.55 indels/100 kbp.

**Figure 4.**
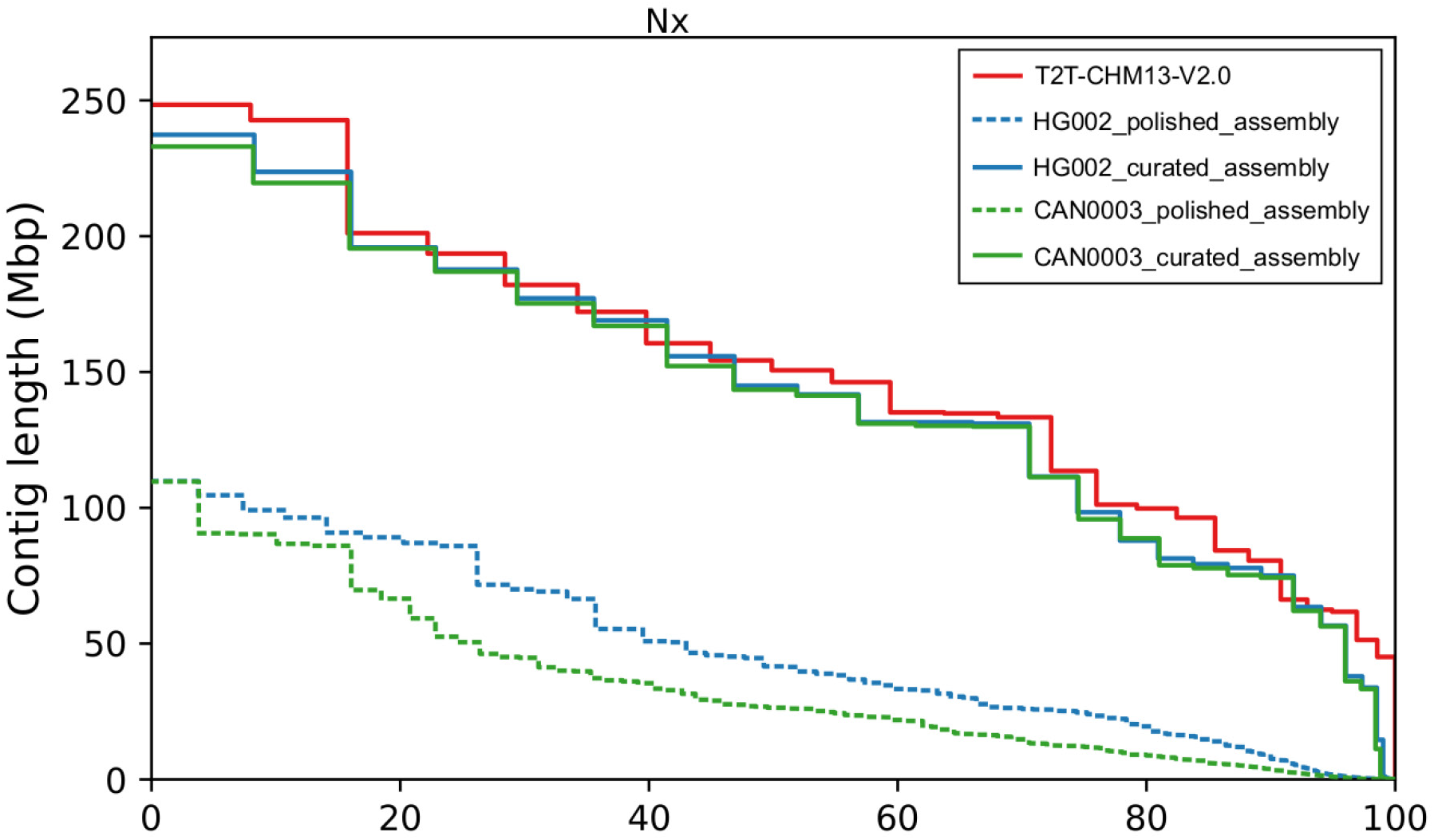
Nx plot of HG002 and CAN0003 polished and curated contigs resulting from *de novo* genome assembly using the Corrected_Flye pipeline and Racon_Pilon polishing schema, as reported by QUAST using T2T-CHM13v2.0 as the reference genome.

The final curated assembly of CAN0003 had 25 scaffolds (1-22 autosomes, X and Y sexual chromosomes, and the mitogenome), plus 395 unplaced contigs. The total length of the assembly was 2,805,379,785 bp, not considering the unplaced contigs, and including 395 gaps summing up to a total gap length of 5,181,142 bp. A comparison of chromosome lengths and gaps between HG002 and CAN0003 genome assemblies, and the T2T-CHM13v2.0 reference is shown in **Supplementary Table 7**.

## Discussion

Given its unique capability to generate very long reads, nanopore sequencing data significantly influences the outcomes of *de novo* genome assembly compared to other sequencing technologies (Lu, Giordano, and Ning 2016; Senol Cali et al. 2019; Nurk et al. 2022). Despite the challenges posed by the higher per base error rates compared to other technologies (H. Zhang, Jain, and Aluru 2019), this benchmarking study highlights the potential of nanopore sequencing to span large genomic regions, contributing to enhance the contiguity in the results. For the first time, we compared up to 11 different pipelines for *de novo* genome assembly of human whole-genomes using nanopore reads from a reference GIAB sample, and up to four different polishing combination schemes. We found that the best performing combination, although not in terms of the required computational time and resources, was to filter out nanopore reads for a minimal size (<1,000 bp), correct them with short reads, and to run Flye assembler, followed by two rounds of Racon and one run of Pilon to polish the sequence with short reads. We then validated the results in sequencing datasets obtained from a routine laboratory sample, and therefore with suboptimal DNA integrity due to handling and storage conditions. The latter comparison provided a comparable number of scaffolds and a total length for the nuclear chromosomes, although having more unplaced contigs and sequence gaps than the reference materials. Finally, we provide a Nextflow-based implementation of the complete pipeline providing this best performing option for *de novo* genome assembly to enable parallelization and dependency management by any user.

Diverse studies have provided comprehensive views of the advantages and limitations of the sequencing technologies and assembly methodologies across different datasets. For instance, Wick and Holt (Wick and Holt 2019) found Flye as a highly reliable assembler on the basis of a benchmarking of eight long-read assemblers using prokaryote samples, although without assessing error-correction and polishing steps. Cosma *et al*. (Cosma et al. 2022) reviewed several long-read *de novo* genome assemblers for ONT, PacBio CLR, and PacBio HiFi reads on diverse eukaryotic genomes using simulated and real data, also concluding that Flye was the best performing tool for ONT and PacBio CLR datasets. A recent study, focusing on the benefits of the multiple sequencing platforms instead of a benchmarking of the software tools, assembled two human samples from the GIAB Consortium comparing multiple sequencing datasets (PacBio CLR and HiFi, ONT, and Illumina short-reads), five *de novo* genome assemblers, and polishing strategies using either only long reads or using both short and long reads (Wang et al. 2023). They concluded that PacBio HiFi was the best technology for genome assembly due to their high base quality, although Flye followed by polishing steps was recommended to assemble ONT reads. The conclusions of our results are consistent with these previous studies, by presenting Flye as an assembly tool with high reliability for ONT data. However, it is worth mentioning that the pursuit of an optimal long-read assembler is shaped by factors such as the sequencing technology and genome complexity. Furthermore, we validated the results across diverse datasets, using not only one of the reference materials from GIAB but also from a routine laboratory sample, underscoring the reproducibility and optimal reliability of adopting a hybrid strategy based on different sequencing technologies (Nurk et al. 2022; Díaz-de Usera et al. 2022).

The polishing schemes that were evaluated in this study relied on Racon, Medaka, and Pilon, three state-of-the-art sequence polishing tools (Senol Cali et al. 2019; Lee et al. 2021). These results demonstrate the importance of polishing draft assemblies for the construction of high-quality reference genomes by improving accuracy, assembly gaps, and potential assembly errors and misassemblies, as reported by recent studies. Chen *et al. (Z. Chen, Erickson, and Meng 2021)* studied the impact of polishing ONT-based bacterial assemblies with Illumina short-reads using two polishing tools. They concluded that NextPolish and, at least, two rounds of Pilon result in similar accuracy levels. As a particular case, Mc Cartney *et al*. (Mc Cartney et al. 2021) used Illumina and PacBio HiFi reads to apply accurate assembly corrections on the T2T-CHM13v0.9 human genome assembly, aiming to improve consensus accuracy, filling gaps, and fixing misassemblies (Fang and Wang 2022). Here, we opted to provide a comparison of various polishing tools and combinations, revealing that a Racon_Pilon scheme achieved a balanced improvement in contiguity and accuracy.

There are some limitations of this study. We have used data from a specific ONT basecalling and a particular PromethION flow cell version, but it is known that the development in this field by ONT is in continuous and rapid improvement (Rang, Kloosterman, and de Ridder 2018; Sereika et al. 2022). Therefore, future studies using more accurate ONT basecallers could lead to significant improvements in genome assemblies. Additionally, in recent years, ONT protocols and reagents have been enhanced, enabling the production of ultra-long reads (Jain et al. 2018) and duplex reads (Koren et al. 2024). These advancements are expected to improve accuracy and may impact the computational time necessities, which can make the use of error-correction or polishing steps using short-reads unnecessary (Nie et al. 2024). The benchmarking of the assembly and polishing tools was primarily based on the HG002 reference sample, and while efforts were made to validate the best performing pipeline on a routine laboratory sample (CAN0003), differences in sample quality, DNA purity, and integrity could influence assembly results in other settings. Our approach is primarily focused on the benefits of combining ONT and Illumina technologies, providing a comprehensive evaluation of *de novo* genome assembly strategies. However, it is important to note that the inclusion of additional technologies, such as PacBio (Wenger et al. 2019), optical mapping by Bionano Genomics (Leinonen and Salmela 2020), or other techniques such as Hi-C to leverage proximity regions to inform the assembly (van Berkum et al. 2010), could further enrich the diversity of genomic data and potentially enhance the overall quality of assembly outcomes (Kim et al. 2019; Ghurye and Pop 2019; Takayama et al. 2021). Some of these other technologies offer unique advantages, such as higher base-level accuracy, improvements in the detection of structural variants, or refinements in phasing and scaffolding, and their integration into future studies could contribute to a more nuanced understanding of the individual differences in the genomic landscapes and the impact in disease. Our study, while insightful within the scope of ONT and Illumina, prompts future investigations to explore the synergies and optimizations by incorporating a broader spectrum of sequencing technologies.

The findings of this study underscore the benefit of integrating long-read sequencing, particularly from ONT platforms (Jain et al. 2018), with short-reads for hybrid *de novo* genome assembly of human whole genomes. Utilizing a combination of base-level error-correction tools, such as Ratatosk, and advanced assembly pipelines and polishers allowed us to obtain assemblies with high accuracy and completeness. Still, the observed differences in processing times and memory utilization among the tested pipelines emphasize the importance of selecting bioinformatics tools that adapt to the available computational resources and project timelines, especially in large-scale genomic studies. Our findings contribute valuable insights and guidance for researchers navigating the complexities of *de novo* genome assembly in diverse genomic contexts. Continuous algorithmic development, scalability optimization, and standardization efforts are essential for the evolving landscape of genomic studies, ensuring the adaptability and reliability of bioinformatics tools in deciphering complex genomes with unprecedented precision.

## Code and data availability

https://github.com/genomicsITER/hybridassembly

## Ethics statement

The study was approved by the Research Ethics Committee of the Hospital Universitario Nuestra Señora de Candelaria (CHUNSC_2020_95) and performed according to The Code of Ethics of the World Medical Association (Declaration of Helsinki).

## Acknowledgements

We would like to thank the support from our colleagues from the Teide-HPC Supercomputing facility (http://teidehpc.iter.es/en), which was funded by INP-2011-0063-PCT-430000-ACT (INNPLANTA program) from the Spanish Ministry of Economy and Competitiveness. AMB, LARR and JMLS acknowledge the training support provided by the University of La Laguna.

## Supplementary Material

### Comprehensive Score (CS) calculation

Several metrics from different bioinformatics tools were integrated in the CS calculation to comprehensively evaluate the quality of the assembled genomes provided by the assemblers and pipelines, similar to what has been described elsewhere (Zhang et al. 2022). The CS score integrates metrics provided by: 1) QUAST: including contig numbers (*contigs*), N50 length in Mb (*N50*), the number of mismatches (*mismatches*), and indels per 100kb (*indels*); 2) BUSCO: including the number of complete genes evaluated (*completeness*); and 3) Merqury: in particular the consensus quality value (*QV*) taking advantage of the *k-mer* based assembly evaluation using Illumina reads. These metrics were integrated into the following equation (1):

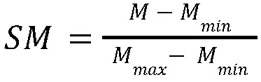

where each metric (*M*) was scaled to [0, 1] by Min-Max normalization across different assembly or polishing pipelines and used to derive a Scaled Metric (*SM*). The *M_min_* or *M_max_* correspond to the minimum or maximum value of *M* among all results to be evaluated. Since high-quality assemblies are expected to have high *SM* in *N50*, *completeness*, and *QV* and low *SM* for *contigs*, *mismatches*, and *indels*, the following equation (2):

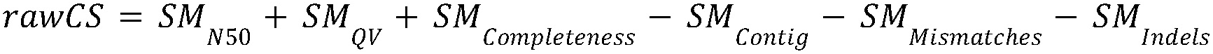

defines the Raw Comprehensive Score (*rawCS*) by summing across the six *SM* values, whose coefficients were set as 1 for the former three metrics and -1 for the latter three metrics to integrate the positive and negative contribution of each. Finally, to obtain the CS, the rawCS was rescaled to [0, 1] by Mix-Max normalization using the following equation (3):

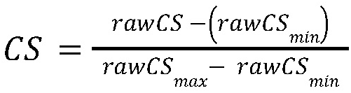

where *rawCS_min_* and *rawCS_max_* correspond to the minimum and maximum theoretical value of the *rawCS*, which can take the values of -3 and 3, respectively.

**Supplementary Table 1.** Quality Control results of genomic DNA after purification of the validation sample.

| Sample | TapeStation size (Kb) | Qubit ng/uL | NanoDrop ng/uL | NanoDrop 260/280 | NanoDrop 260/230 |
| --- | --- | --- | --- | --- | --- |
| CAN0003 | 48 | 36 | 33.97 | 1.83 | 2.55 |

**Supplementary Table 2.** Complete HG002 assembly results. T2T-CHM13v2.0 genome was used as reference in QUAST evaluation. Best value of each metric is highlighted.

| Input reads | Assembly type | Assembler | QUAST |  |  |  |  |  |  |  |  |  | Merqury |  | BUSCO* |  |  |  |  |
| --- | --- | --- | --- | --- | --- | --- | --- | --- | --- | --- | --- | --- | --- | --- | --- | --- | --- | --- | --- |
|  |  |  | Total length (Gbp) | # contigs | N50 (Mbp) | L50 | Largest contig (Mbp) | Genome fraction (%) | NA50 (Mbp) | # misassemblies | # mismatches / 100 kbp | # indels / 100 kbp | k-mer completeness (%) | Quality value | C | S | D | F | M |
| Filtered | LR only assemblies | Shasta | 2.91 | 2,167 | 41.22 | **20** | **138.11** | 91.89 | 30.47 | 4,413 | 191.76 | 106.98 | 96.24 | 32.70 | 12,258 | 12,049 | 209 | 463 | 1,059 |
|  |  | Flye | 2.86 | 845 | 38.29 | 23 | 108.41 | 91.25 | 31.42 | 925 | 141.24 | 107.35 | 96.18 | 33.73 | 12,249 | 12,024 | 225 | 552 | 979 |
|  |  | Raven | 2.86 | **313** | 32.50 | 28 | 109.44 | 90.61 | 26.09 | 907 | 151.74 | 115.62 | 95.66 | 32.57 | 12,133 | 11,926 | 207 | 566 | 1,081 |
|  |  | wtdbg2 | 2.93 | 8,712 | 10.34 | 78 | 44.81 | 86.54 | 6.97 | 1,986 | 228.31 | 274.81 | 90.07 | 25.82 | 10,849 | 10,666 | 183 | 726 | 2,205 |
| Filtered and corrected | LR only assemblies | Shasta | 2.87 | 1,153 | 8.75 | 104 | 47.52 | 91.56 | 8.05 | 426 | 115.90 | 41.65 | 96.85 | 40.69 | 12,974 | 12,747 | 227 | 214 | 592 |
|  |  | Flye | 2.91 | 819 | **41.59** | 21 | 109.82 | **92.41** | **33.17** | 3,136 | 127.48 | 26.16 | **97.50** | **47.84** | **13,196** | **12,969** | 227 | **145** | 439 |
|  |  | Raven | 2.85 | 583 | 28.21 | 33 | 83.95 | 86.33 | 0.49 | 137 | **97.40** | 32.18 | 96.66 | 40.54 | 13,176 | 12,937 | 239 | 149 | 455 |
|  |  | wtdbg2 | **3.28** | 18,052 | 3.74 | 209 | 28.23 | 87.53 | 3.08 | 3,854 | 255.29 | 62.51 | 93.54 | 30.95 | 13,176 | 11,915 | 671 | 205 | 989 |
| Filtered | Hybrid assemblies | WENGAN | 2.83 | 1,314 | 29.31 | 32 | 109.74 | 90.56 | 23.76 | 480 | 111.85 | 26.87 | 97.21 | 45.32 | 13,164 | 12,957 | 207 | 148 | 468 |
|  |  | MaSuRCA | 2.85 | 792 | 16.66 | 48 | 88.12 | 90.98 | 13.19 | 916 | 136.12 | **25.41** | 97.27 | 43.95 | 13,186 | 12,966 | 220 | 157 | **437** |
|  |  | HASLR | 2.73 | 6,030 | 1.03 | 752 | 7.53 | 87.38 | 1.02 | **113** | 112.74 | 70.57 | 95.38 | 36.26 | 12,620 | 12,455 | **165** | 368 | 792 |

C: Complete BUSCOs, S: Complete and single-copy BUSCOs, D: Complete and duplicated BUSCOs, F: Fragmented BUSCOs, M: Missing BUSCOs
\*Total BUSCO groups searched: 13,780

**Supplementary Table 3.** CS calculations of HG002 assembly results. Best CS value is highlighted.

| Input reads | Assembly type | Assembler | # contigs | N50 (Mbp) | Mismatches | Indels | QV | Completeness | SM<sub>contig</sub> | SM<sub>N50</sub> | SM<sub>mismatches</sub> | SM<sub>indels</sub> | SM<sub>QV</sub> | SM<sub>completeness</sub> | raw_CS | CS |
| --- | --- | --- | --- | --- | --- | --- | --- | --- | --- | --- | --- | --- | --- | --- | --- | --- |
| Filtered | LR only assemblies | Shasta | 2,167 | 41.22 | 191.76 | 106.98 | 32.70 | 12,258 | 0.10 | 0.99 | 0.60 | 0.33 | 0.31 | 0.60 | 0.87 | 0.65 |
|  |  | Flye | 845 | 38.29 | 141.24 | 107.35 | 33.73 | 12,249 | 0.03 | 0.92 | 0.28 | 0.33 | 0.36 | 0.60 | 1.24 | 0.71 |
|  |  | Raven | 313 | 32.50 | 151.74 | 115.62 | 32.57 | 12,133 | 0.00 | 0.78 | 0.34 | 0.36 | 0.31 | 0.55 | 0.92 | 0.65 |
|  |  | wtdbg2 | 8,712 | 10.34 | 228.31 | 274.81 | 25.82 | 10,849 | 0.47 | 0.23 | 0.83 | 1.00 | 0.00 | 0.00 | -2.07 | 0.15 |
| Filtered and corrected | LR only assemblies | Shasta | 1,153 | 8.75 | 115.90 | 41.65 | 40.69 | 12,974 | 0.05 | 0.19 | 0.12 | 0.07 | 0.68 | 0.91 | 1.54 | 0.76 |
|  |  | Flye | 819 | 41.59 | 127.48 | 26.16 | 47.84 | 13,196 | 0.03 | 1.00 | 0.19 | 0.00 | 1.00 | 1.00 | 2.78 | **0.96** |
|  |  | Raven | 583 | 28.21 | 97.40 | 32.18 | 40.54 | 13,176 | 0.02 | 0.67 | 0.00 | 0.03 | 0.67 | 0.99 | 2.29 | 0.88 |
|  |  | wtdbg2 | 18,052 | 3.74 | 255.29 | 62.51 | 30.95 | 12,586 | 1.00 | 0.07 | 1.00 | 0.15 | 0.23 | 0.74 | -1.11 | 0.32 |
| Filtered | Hybrid assemblies | WENGAN | 1,314 | 29.31 | 111.85 | 26.87 | 45.32 | 13,164 | 0.06 | 0.70 | 0.09 | 0.01 | 0.89 | 0.99 | 2.42 | 0.90 |
|  |  | MaSuRCA | 792 | 16.66 | 136.12 | 25.41 | 43.95 | 13,186 | 0.03 | 0.39 | 0.25 | 0.00 | 0.82 | 1.00 | 1.93 | 0.82 |
|  |  | HASLR | 6,030 | 1.03 | 112.74 | 70.57 | 36.26 | 12,620 | 0.32 | 0.00 | 0.10 | 0.18 | 0.47 | 0.75 | 0.63 | 0.60 |

\*SM: Scaled metric

**Supplementary Table 4.** Complete HG002 polishing results for the best assembly pipeline. T2T-CHM13v2.0 genome was used as reference in QUAST evaluation. Best value of each metric is highlighted.

| Input reads | Assembler | Polishing pipeline | QUAST |  |  |  |  |  |  |  |  |  | Merqury |  | BUSCO* |  |  |  |  |
| --- | --- | --- | --- | --- | --- | --- | --- | --- | --- | --- | --- | --- | --- | --- | --- | --- | --- | --- | --- |
|  |  |  | Total length (Gbp) | Total contigs | N50 (Mbp) | L50 | Largest contig (Mbp) | Genome fraction (%) | NA50 (Mbp) | # misassemblies | # mismatches / 100 kbp | # indels / 100 kbp | k-mer completeness (%) | Quality value | C | S | D | F | M |
| Filtered and corrected | Flye | Racon_Pilon | 2.91 | **800** | 41.59 | **21** | 109.83 | 92.30 | 31.63 | **3,210** | 131.39 | 24.53 | 97.49 | **46.58** | 13,198 | 12,971 | 227 | 147 | **435** |
|  |  | Racon_Medaka | **2.92** | **800** | **41.62** | **21** | **109.91** | 92.27 | 31.53 | 3,264 | 133.99 | 28.02 | 97.47 | 42.68 | 13,196 | 12,969 | 227 | 146 | 438 |
|  |  | Medaka_Pilon | 2.91 | 819 | 41.59 | **21** | 109.84 | **92.38** | **33.28** | 3,343 | **125.82** | **24.21** | **97.53** | 45.77 | **13,199** | **12,973** | 226 | **145** | 436 |
|  |  | Racon_Medaka_Pilon | 2.91 | **800** | 41.60 | **21** | 109.87 | 92.30 | 33.18 | 3,239 | 127.93 | 24.57 | 97.51 | 44.21 | 13,196 | 12,969 | 227 | **145** | 439 |

C: Complete BUSCOs, S: Complete and single-copy BUSCOs, D: Complete and duplicated BUSCOs, F: Fragmented BUSCOs, M: Missing BUSCOs
\*Total BUSCO groups searched: 13,780

**Supplementary Table 5.**
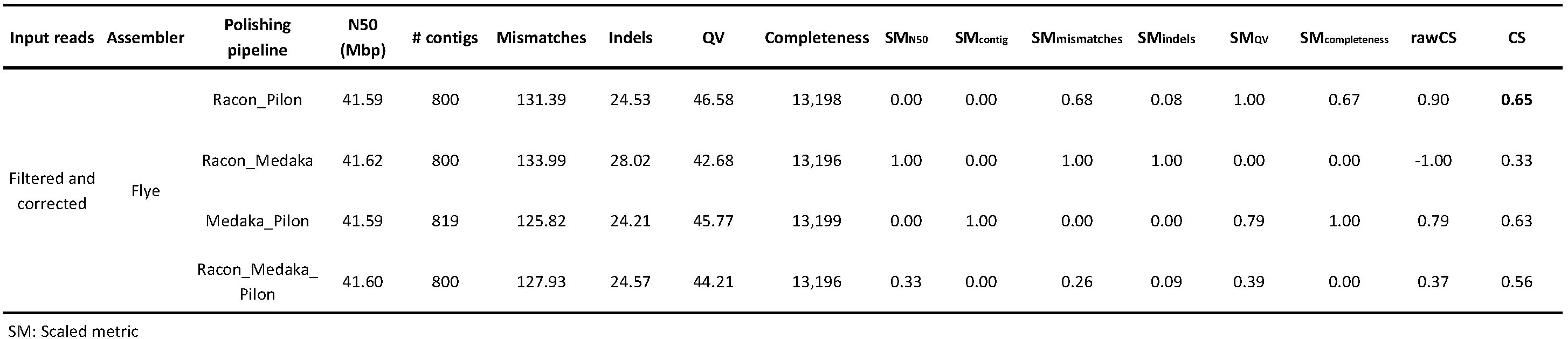
CS calculations of HG002 polishing results. Best CS value is highlighted.

| Input reads | Assembler | Polishing pipeline | N50 (Mbp) | # contigs | Mismatches | Indels | QV | Completeness | SM<sub>N50</sub> | SM<sub>contig</sub> | SM<sub>mismatches</sub> | SM<sub>indels</sub> | SM<sub>QV</sub> | SM<sub>completeness</sub> | rawCS | CS |
| --- | --- | --- | --- | --- | --- | --- | --- | --- | --- | --- | --- | --- | --- | --- | --- | --- |
| Filtered and corrected | Flye | Racon_Pilon | 41.59 | 800 | 131.39 | 24.53 | 46.58 | 13,198 | 0.00 | 0.00 | 0.68 | 0.08 | 1.00 | 0.67 | 0.90 | **0.65** |
|  |  | Racon_Medaka | 41.62 | 800 | 133.99 | 28.02 | 42.68 | 13,196 | 1.00 | 0.00 | 1.00 | 1.00 | 0.00 | 0.00 | -1.00 | 0.33 |
|  |  | Medaka_Pilon | 41.59 | 819 | 125.82 | 24.21 | 45.77 | 13,199 | 0.00 | 1.00 | 0.00 | 0.00 | 0.79 | 1.00 | 0.79 | 0.63 |
|  |  | Racon_Medaka_Pilon | 41.60 | 800 | 127.93 | 24.57 | 44.21 | 13,196 | 0.33 | 0.00 | 0.26 | 0.09 | 0.39 | 0.00 | 0.37 | 0.56 |

SM: Scaled metric

**Supplementary Table 6.** Complete HG002 results of each curation step. T2T-CHM13v2.0 genome was used as reference in QUAST evaluation.

| Input reads | Assembler | Polishing pipeline | Curation step | Tools | QUAST |  |  |  |  |  |  |  |  |  | Merqury |  | BUSCO* |  |  |  |  |  |
| --- | --- | --- | --- | --- | --- | --- | --- | --- | --- | --- | --- | --- | --- | --- | --- | --- | --- | --- | --- | --- | --- | --- |
|  |  |  |  |  | Total length (Gbp) | Total contigs | N50 (Mbp) | L50 | Largest contig (Mbp) | Genome fraction (%) | NA50 (Mbp) | # misassemblies | # N's per 100 kbp | # mismatches / 100 kbp | # indels / 100 kbp | k-mer completeness (%) | Quality value | C | S | D | F | M |
| Filtered and corrected | Flye | Racon_Pilon | Purge haplotigs and overlaps | purge_dups | 2.87 | 409 | 41.59 | 21 | 109.83 | 91.54 | 31.63 | 1,286 | 0.00 | 115.85 | 23.90 | 97.32 | 47.94 | 13,225 | 13,001 | 224 | 124 | 431 |
|  |  |  | Misassemblies correction | RagTag | 2.87 | 518 | 41.39 | 22 | 109.83 | 91.55 | 31.63 | 1,342 | 0.00 | 116.27 | 23.92 | 97.32 | 47.94 | 13,225 | 13,000 | 225 | 125 | 431 |
|  |  |  | Scaffolding | RagTag | 2.87 | 230 | 145.01 | 8 | 237.34 | 91.54 | 50.04 | 1,546 | 76.28 | 118.06 | 23.87 | 97.32 | 47.94 | 13,221 | 13,006 | 215 | 125 | 434 |
|  |  |  | Close gaps | TGS-GapCloser | 2.87 | 230 | 144.97 | 8 | 237.34 | 91.58 | 44.66 | 1,571 | 38.03 | 118.94 | 23.94 | 97.32 | 47.65 | 13,221 | 13,005 | 216 | 125 | 434 |

C: Complete BUSCOs, S: Complete and single-copy BUSCOs, D: Complete and duplicated BUSCOs, F: Fragmented BUSCOs, M: Missing BUSCOs
\*Total BUSCO groups searched: 13,780

**Supplementary Table 7.** Comparison of chromosome lengths and gaps between the T2T-CHM13v2.0 reference and HG002 and CAN0003 curated assemblies.

| Chromosome | T2T-CHM13v2.0 |  | HG002 |  | CAN0003 |  |  |
| --- | --- | --- | --- | --- | --- | --- | --- |
|  | Length (bp) | Length (bp) | Gap length (bp) | # of gaps | Length (bp) | Gap length (bp) | # of gaps |
| 1 | 248,387,328 | 223,695,053 | 157,510 | 19 | 219,557,404 | 359,976 | 32 |
| 2 | 242,696,752 | 237,337,981 | 146,350 | 14 | 232,949,127 | 382,875 | 32 |
| 3 | 201,105,948 | 195,886,542 | 47,472 | 6 | 195,428,087 | 39,731 | 6 |
| 4 | 193,574,945 | 187,717,499 | 40,047 | 5 | 186,937,478 | 166,621 | 13 |
| 5 | 182,045,439 | 177,128,092 | 27,784 | 6 | 175,229,402 | 86,823 | 21 |
| 6 | 172,126,628 | 169,003,213 | 500 | 5 | 166,952,613 | 88,983 | 6 |
| 7 | 160,567,428 | 155,774,717 | 99,511 | 12 | 152,151,974 | 489,452 | 31 |
| 8 | 146,259,331 | 141,744,978 | 400 | 4 | 141,244,837 | 78,807 | 6 |
| 9 | 150,617,247 | 111,517,537 | 68,337 | 7 | 111,218,482 | 343,197 | 35 |
| 10 | 134,758,134 | 131,533,104 | 600 | 6 | 129,873,681 | 327,859 | 18 |
| 11 | 135,127,769 | 131,497,776 | 100 | 1 | 130,123,708 | 93,024 | 5 |
| 12 | 133,324,548 | 130,984,586 | 100 | 1 | 130,941,057 | 87,014 | 5 |
| 13 | 113,566,686 | 98,374,707 | 1,100 | 11 | 95,758,623 | 120,241 | 9 |
| 14 | 101,161,492 | 87,905,942 | 559 | 4 | 88,708,172 | 79,159 | 9 |
| 15 | 99,753,195 | 81,334,459 | 1,200 | 12 | 78,793,148 | 340,367 | 27 |
| 16 | 96,330,374 | 77,844,360 | 66,759 | 15 | 75,215,631 | 205,789 | 20 |
| 17 | 84,276,897 | 79,284,333 | 186,075 | 11 | 77,713,116 | 650,353 | 27 |
| 18 | 80,542,538 | 75,051,009 | 200 | 2 | 74,252,416 | 47,455 | 8 |
| 19 | 61,707,364 | 56,615,862 | 32,246 | 3 | 56,309,012 | 135,063 | 10 |
| 20 | 66,210,255 | 63,485,542 | 600 | 6 | 62,013,617 | 102,861 | 8 |
| 21 | 45,090,682 | 33,793,004 | 200 | 2 | 33,235,988 | 500 | 5 |
| 22 | 51,324,926 | 37,964,771 | 13,909 | 14 | 36,124,491 | 175,333 | 14 |
| X | 154,259,566 | 144,968,395 | 200,271 | 9 | 143,459,348 | 619,970 | 31 |
| Y | 62,460,029 | 14,560,519 | 800 | 8 | 11,171,801 | 159,689 | 17 |
| MT | 16,569 | 16,571 | 0 | 0 | 16,572 | 0 | 0 |
| Total | 3,117,292,070 | 2,845,020,552 | 1,092,630 | 183 | 2,805,379,785 | 5,181,142 | 395 |

**Supplementary Table 8.** Complete CAN0003 results of each pipeline step. T2T-CHM13v2.0 genome was used as reference in QUAST evaluation.

| Input reads | Pipeline step | Tools | QUAST |  |  |  |  |  |  |  |  |  | Merqury |  | BUSCO* |  |  |  |  |  |
| --- | --- | --- | --- | --- | --- | --- | --- | --- | --- | --- | --- | --- | --- | --- | --- | --- | --- | --- | --- | --- |
|  |  |  | Total length (Gbp) | Total contigs | N50 (Mbp) | L50 | Largest contig (Mbp) | Genome fraction (%) | NA50 (Mbp) | # misassemblies | # N's per 100 kbp | # mismatches / 100 kbp | # indels / 100 kbp | k-mer completeness (%) | Quality value | C | S | D | F | M |
| Filtered and corrected | Assembly | Flye | 2.89 | 1,792 | 26.37 | 30 | 109.62 | 91.72 | 23.52 | 2,614 | 0.00 | 124.82 | 35.42 | 97.40 | 43.06 | 13,172 | 12,950 | 222 | 161 | 447 |
|  | Polishing pipeline | Racon_Pilon | 2.89 | 1,738 | 26.38 | 30 | 109.64 | 91.53 | 22.30 | 2,797 | 0.00 | 163.02 | 31.30 | 97.42 | 42.46 | 13,182 | 12,960 | 222 | 156 | 442 |
|  | Purge haplotigs and overlaps | purge_dups | 2.83 | 860 | 26.76 | 29 | 109.64 | 90.40 | 22.86 | 1,074 | 0.00 | 147.71 | 30.59 | 97.06 | 42.94 | 13,175 | 12,964 | 211 | 159 | 446 |
|  | Misassemblies correction | RagTag | 2.83 | 913 | 26.76 | 29 | 109.64 | 90.40 | 22.86 | 1,073 | 0.00 | 147.79 | 30.59 | 97.06 | 42.94 | 13,175 | 12,964 | 211 | 159 | 446 |
|  | Scaffolding | RagTag | 2.84 | 419 | 143.53 | 8 | 232.95 | 90.38 | 33.18 | 1,358 | 211.09 | 151.40 | 30.53 | 97.06 | 42.94 | 13,179 | 12,983 | 196 | 156 | 445 |
|  | Close gaps | TGS-GapCloser | 2.84 | 419 | 143.46 | 8 | 232.95 | 90.39 | 32.77 | 1,362 | 182.54 | 151.72 | 30.55 | 97.06 | 42.92 | 13,179 | 12,982 | 197 | 156 | 445 |

C: Complete BUSCOs, S: Complete and single-copy BUSCOs, D: Complete and duplicated BUSCOs, F: Fragmented BUSCOs, M: Missing BUSCOs
\*Total BUSCO groups searched: 13,780

